# A Krüppel-like factor establishes cellular heterogeneity during schistosome tegumental maintenance

**DOI:** 10.1101/2024.07.12.603265

**Authors:** Lu Zhao, George R. Wendt, James. J. Collins

**Affiliations:** Department of Pharmacology, University of Texas Southwestern Medical Center, Dallas, TX

## Abstract

Schistosomes are blood-dwelling parasitic flatworms that rely on a syncytial surface coat, known as the tegument, for long-term survival and immune evasion in the blood of their human hosts. Previous studies have shown that cells within the tegumental syncytium are perpetually turned over and renewed by somatic stem cells called neoblasts. Yet, little is known about this renewal process on a molecular level. Here, we characterized a Krüppel-like factor 4 (*klf4*) using a combination of bulk and single cell RNAseq approaches and demonstrate that *klf4* is essential for the maintenance of a specific tegumental lineage, resulting in the loss of a subpopulation of molecularly-unique tegument cells. Thus, *klf4* is critical for maintaining the balance between different tegumental progenitor pools, thereby fine-tuning the molecular composition of the mature tegument. Understanding these distinct tegumental cell populations is expected to provide insights into parasite defense mechanisms and suggest new avenues for therapeutics.

## Introduction

Schistosomes are parasitic flatworms that infect more than 250 million people around the world causing tens of thousands of deaths and devastating morbidity in the developing world^1^. These parasites are capable of surviving inside the blood of their human host for decades in the face of the host’s immune system and the tremendous physical challenges imposed by living in the circulation^2-4^. The parasite’s syncytial skin-like surface coat called the tegument is thought to be essential for not just surviving but thriving in this hostile environment^5-7^. As the interface between the parasite and the host, the tegument is involved in acquiring nutrients to support worm growth^8,9^ and for evading host immune responses^7^.

The tegument has a complicated architecture, comprised of thousands of nucleated cell bodies that connect to the outer tegument via cellular projections extending through the body wall muscles^10,11^. Interestingly, previous studies have shown that schistosome tegumental cell bodies are subject to perpetual turnover and replacement^11,12^. This replacement of tegumental cell bodies is driven by population of somatic stem cells, called neoblasts, that specify a pool of short-lived *tsp-2*^+^ tegument progenitor cells^12^ that migrate through the worm to fuse with the tegumental syncytium^11^. Though the tegument is a continuous syncytium that covers the entire surface of schistosome, meaning all tegument cells ostensibly share a common cytoplasm, we previously uncovered signs of molecular heterogeneity in both the *tsp-2*^+^ tegument progenitor pool as well as the mature tegument itself^11^. Subsequent single cell RNAseq (scRNAseq) studies went on to reveal two distinct tegument associated lineages (TALs): clusters of cells that were molecularly unique from one another yet both expressed genes associated with the tegument^13^. Thus, important questions remain about the purpose of these two TALs. In particular, whether both TALs represent true tegumental cell progenitor populations and, if so, whether they play a role in establishing the molecular heterogeneity of the tegument.

Here, we examine a Krüppel-like factor 4 (KLF4) homolog that was specifically enriched in one of the two TALs. Knockdown of *klf4* resulted in ablation of the entire *klf4*^+^ TAL and resulted in a concomitant loss of a molecularly-unique subpopulation of tegument cells. Surprisingly, loss of the *klf4*^+^ TAL had no effect on the overall maintenance of tegument cell numbers. Instead, loss of the *klf4*^+^ TAL was accompanied by a compensatory increase in the flux through the intact TAL. This suggests that parasites rely upon the balance between two different tegument progenitor lineages to fine-tune the molecular identity of the mature tegument.

## Results

### Krüppel-like factor (*klf4)* is enriched in tegument associated lineage of schistosome

Our recently published scRNAseq atlas of adult worms^13^, identified two lineages predicted to produce tegumental cells (**Fig. 1a**) based on their expression of the mRNAs that encode the tegumental protein TSP-2^14^ **(Fig. 1b);** for brevity, we will refer to these as tegument associated lineages (TALs). One TAL was characterized by the expression of *tsp-2* (*Smp_335630*) and *sm13* (*Smp_195190*) **(Fig. 1c)**, whereas the other TAL expressed *tsp-2* along with *Endoglycoceramidase* (*egc, Smp_314170*), *meg-1* (*Smp_122630*), and *zfp-1-1*^11^ **(Fig. 1d-f)**. To explore the functions of these lineages, we explored our scRNAseq data and found that a homolog of a Krüppel-like factor 4 (KLF4) transcription factor-encoding gene (*Smp_018170, klf4* for brevity) (**Supplementary Fig. 1a**) was highly expressed in the *egc, meg-1*, and *zfp-1-1* TAL (**Fig. 2a**). Whole mount *in situ* hybridization (WISH) showed that this gene is expressed in discrete cells throughout the body (**Fig. 2b**) and double fluorescence *in-situ* hybridization (FISH) confirmed the expression of *klf4* in *egc*^+^, *meg-1*^+^ and *zfp-1-1*^+^ cells (**Fig. 2c**). Additionally, *klf4* was co-expressed with the tegument progenitor marker *tsp-2* (**Fig. 2c bottom**), but not in cells expressing *sm13* that mark the other TAL lineage (**Supplementary Fig. 1b**). These data suggest *klf4* is abundantly and specifically expressed in one of the two putative TALs.

**Fig. 1:**
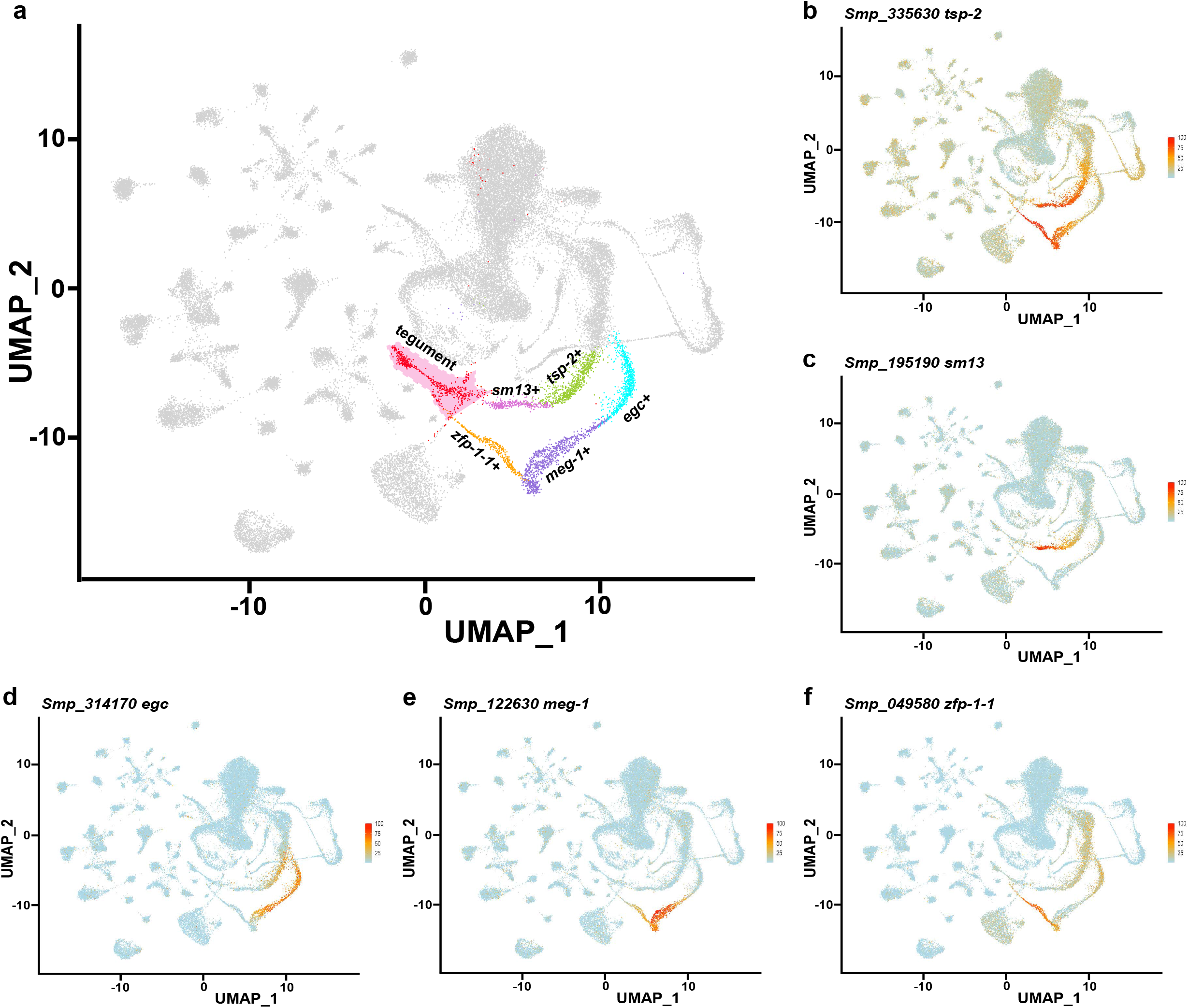
scRNAseq atlas of the adult schistosome reveals two tegument associated lineages (TALs). **a** Uniform Manifold Approximation and Projection (UMAP) showing a scRNAseq atlas of the adult schistosome revealing two lineages predicted to produce tegumental cells. One lineage is characterized by the expression of *tsp-2* and *sm13*, the other lineage expresses *tsp-2* along with *egc, meg-1* and *zfp-1-1*. **b-f** UMAP expression of *tsp-2* (**b**), *sm13* (**c**), *egc* (**d**), *meg-1* (**e**) and *zfp-1-1* (**f**) in TALs.

**Fig. 2:**
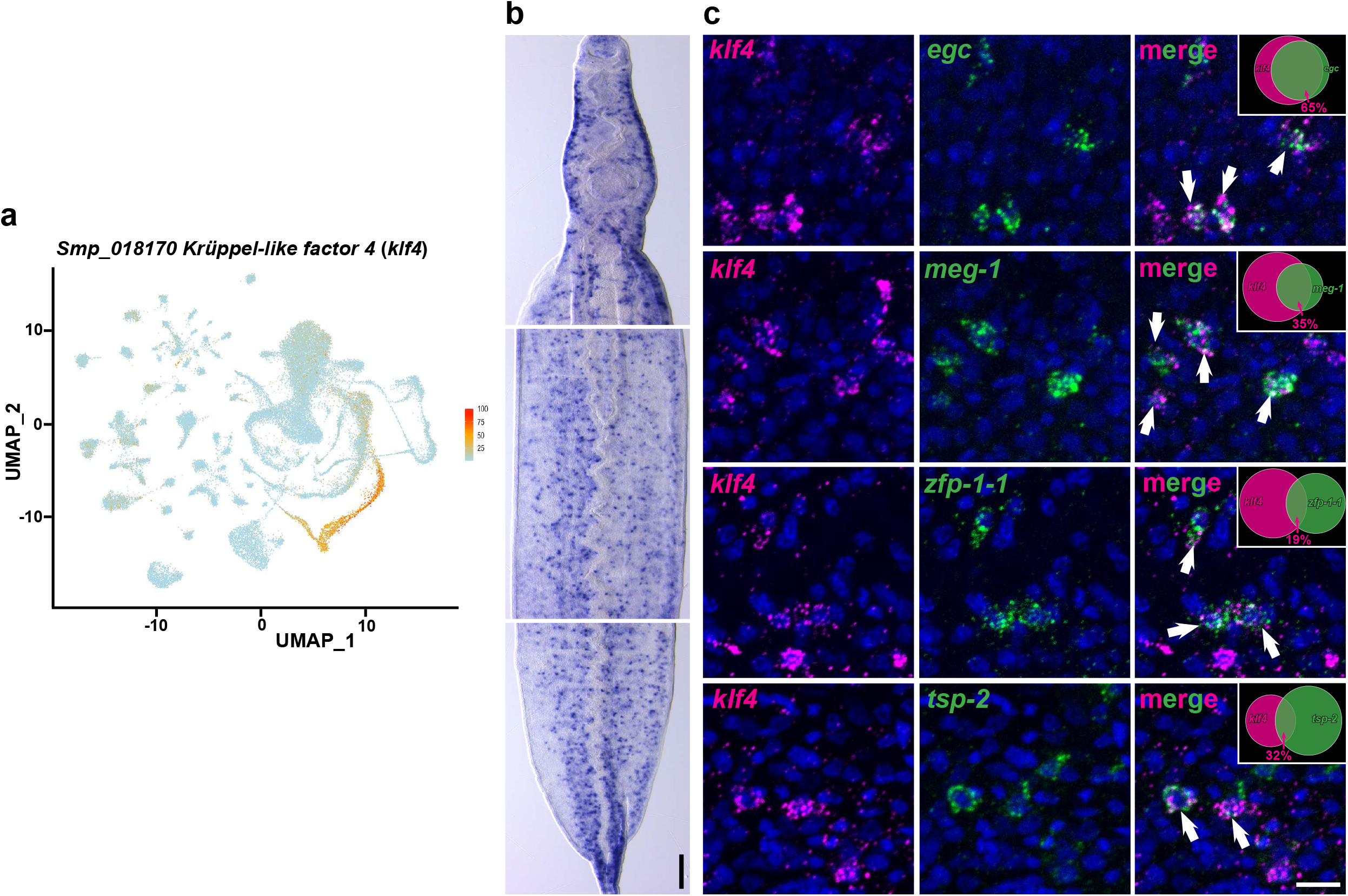
*Klf4* is enriched in schistosome TAL. **a** UMAP showing *klf4* expression in *egc*/*meg-1*/*zfp-1-1* TAL. **b** Colorimetric whole *in-situ* hybridization (WISH) showing expression pattern of *klf4*. Scale bar, 100 µM; anterior towards the top. **c** Double fluorescence *in situ* hybridization (FISH) showing expression of *klf4* relative to the *egc*^+^, *meg-1*^+^, *zfp-1-1*^+^ and *tsp-2*^+^ cells (indicated in arrows). 65%, 35%, 19% and 32% of *klf4*^+^ cells are *egc*^+^ (81/125 cells, n=3 parasites), *meg-1*^+^ (88/255 cells, n=4 parasites), *zfp-1-1*^+^ (59/319 cells, n=6 parasites) and *tsp-2*^+^ (70/217 cells, n=6 parasites), respectively, as indicated in the Venn diagram in the upper-right. Scale bar, 10 µM.

### *klf4* is required for tegument associated lineage (TAL)

To further investigate the function of *klf4*, we performed RNAi of *klf4* on adult male *S. mansoni*, then performed transcriptional profiling using both scRNAseq and bulk RNAseq to characterize cellular and molecular changes associated with *klf4* loss of function (**Fig. 3a**). Our scRNAseq analysis showed that *klf4* RNAi resulted in a complete loss of the *klf4*^+^ TAL (**Fig. 3b and Supplementary Fig. 2a-e; Supplementary Data 1**), and bulk RNAseq analysis revealed 67 differential expressed genes (DEGs), of which 55 are down-regulated genes, following *klf4* RNAi (**Fig 3c, Supplementary Data 2**). Mapping these DEGs onto the scRNAseq atlas, depicted in Fig 3b, demonstrated that 71% (39/55) of the down-regulated genes were enriched in the *klf4*^+^ TAL (**Fig. 3d, Supplementary Fig. 2f**). To confirm these data, we performed FISH and qPCR to evaluate gene expression changes in worms subjected to *klf4* RNAi. The results confirmed a complete loss of *klf4*^+^, *egc*^+^ and *meg-1*^+^ cells, as well as a substantial reduction in *zfp-1-1*^+^ cells (**Fig. 3e, Supplementary Fig. 2g**), without any significant changes in the number of proliferative EdU^+^ neoblasts in these worms (**Fig. 3e, Supplementary Fig. 2h**). Additionally, *klf4* RNAi also led to a ∼15% reduction in *tsp-2* transcript levels (**Fig. 3f-h**), consistent with our bulk RNAseq analysis (**Fig. 3c green dot, Supplementary Data 2**). These results indicate that *klf4* is essential for the maintenance of the *egc*/*meg-1*/*zfp-1-1* TAL.

**Fig. 3:**
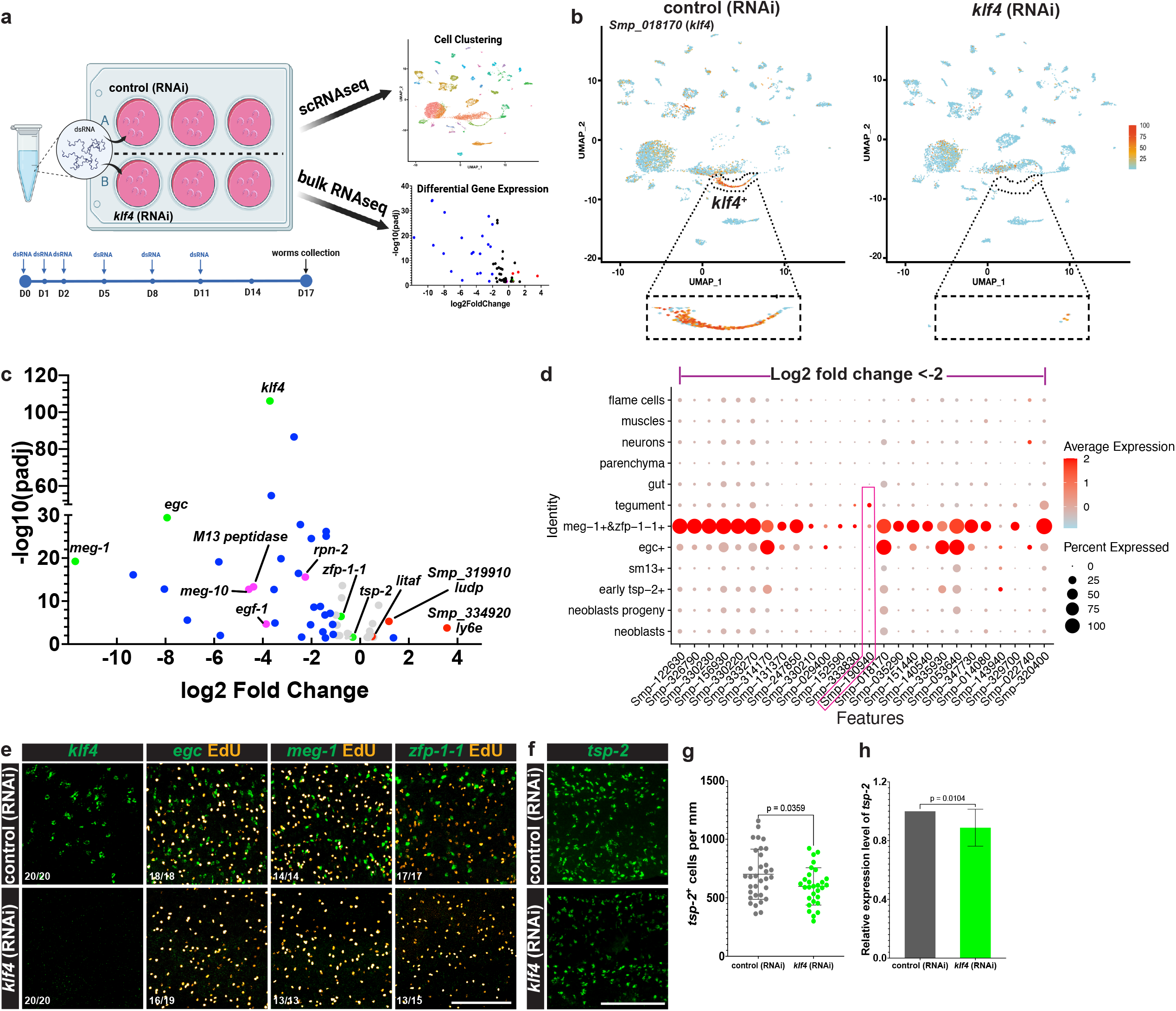
*Klf4* is required for TAL. **a** Graphic depicting the workflow for exploring function of *klf4* by scRNAseq and bulk RNAseq transcriptional profiling analysis. **b** scRNAseq analysis shows *klf4* RNAi resulted in a loss of the *klf4*^+^ TAL. **c** Volcano plot depicting bulk RNAseq analysis. This analysis identified 67 differential expressed genes (*Padj*<0.05, 55 down- and 12 up-regulated genes) following knocking down *klf4*. Grey dots represent genes with a log2 fold change (log2FC) between -1 and 1. Blue dots represent Log2FC<-1 or >1. Green dots indicate marker genes expressed in the *klf4*^+^ TAL. Magenta dots indicate down-regulated genes validated to be expressed in tegument and *zfp-1-1*^+^ cells. Red dots indicate up-regulated genes validated to be expressed in *sm13*^+^ cells. **d** A dot-plot summarizing the expression of the bulk RNAseq down-regulated DEGs (log2 fold change<-2) in clusters from the *klf4* RNAi scRNAseq profile in panel 3B. Cluster populations are on the vertical axis and gene IDs are on the horizontal axis. Expression levels are colored by gene expression (blue = low, red = high). Percentage of cells in the cluster expressing the gene is indicated by the size of the circle. **e** FISH results validating the loss of *klf4*^+^, *egc*^+^, *meg-1*^+^ and a substantial reduction in *zfp-1-1*^+^ cells following *klf4* RNAi. We noted no changes in the number of EdU^+^ proliferative cells (Scale bar, 100 µM). Numbers at bottom left represent the fraction of parasites displaying the observed phenotype. **f** FISH depicting a modest decrease in *tsp-2*^+^ cell number following *klf4* RNAi. Scale bar, 100 µM. **g** Quantification of the number of *tsp-2*^+^ cells per mm of worm. Control (RNAi) n= 33, *klf4* (RNAi) n=29. **h** qPCR quantification of expression of *tsp-2* following *klf4* RNAi. n=12 experiments.

### *Klf4*^+^ TAL is required for producing a specific tegumental subpopulation

Given the profound effects of *klf4* RNAi treatment on the maintenance of the *egc*/*meg-1*/*zfp-1-1* TAL, we evaluated the effects of *klf4* depletion on tegumental maintenance. Surprisingly, despite ablating an entire TAL, knockdown of *klf4* resulted in no change in the total number of tegument cells (**Fig. 4a, 4b**) and likewise had no effect on the transcript levels of definitive tegumental marker *calpain* (**Fig. 4c**). However, our bulk RNAseq studies found that 16% of the down-regulated DEGs (9/55) were expressed in the tegumental cell population identified by scRNAseq (**Supplementary Fig. 2f**). Among them was an *EGF-like domain-containing protein* (*Smp_190940, egf-1*) that was highly downregulated following *klf4* RNAi treatment (**Fig. 3c, 3d, highlighted in pink rectangle**). We observed that *klf4* RNAi resulted in complete loss of cell populations expressing *egf-1* (**Fig. 4d, Supplementary Fig. 3a**). Among these *egf-1*^+^ cells, ∼80% expressed the tegumental marker *calpain* while another ∼20% expressed *zfp-1-1*^+^ (**Fig. 4e top**). These data are consistent with *egf-1* being expressed in a transition state between *zfp-1-1*^*+*^ cells differentiating to definitive *calpain*^*+*^ tegument cells. Similarly, following *klf4* RNAi, we observed a loss of the *meg-10*^*+*^ (*Smp_152590*) cell population (**Fig. 4d bottom**). Similar to *egf-1*, the majority (∼60%) of *meg-10*^+^ cells co-expressed *calpain*^*+*^, suggesting that they are definitive tegument cells (**Fig. 4e bottom**). We noted similar expression patterns with genes encoding an *M13 peptidase* homolog (*Smp_333830*) and a *Ribophorin-2* homolog (*rpn-2, Smp_329700*) (**Supplementary Fig. 3b-d**). Taken together, these data suggest that *klf4* is essential for the maintenance of a subset of tegumental cells that express *egf-1 and meg-10* but is dispensable for the overall maintenance of tegumental cell numbers.

**Fig. 4:**
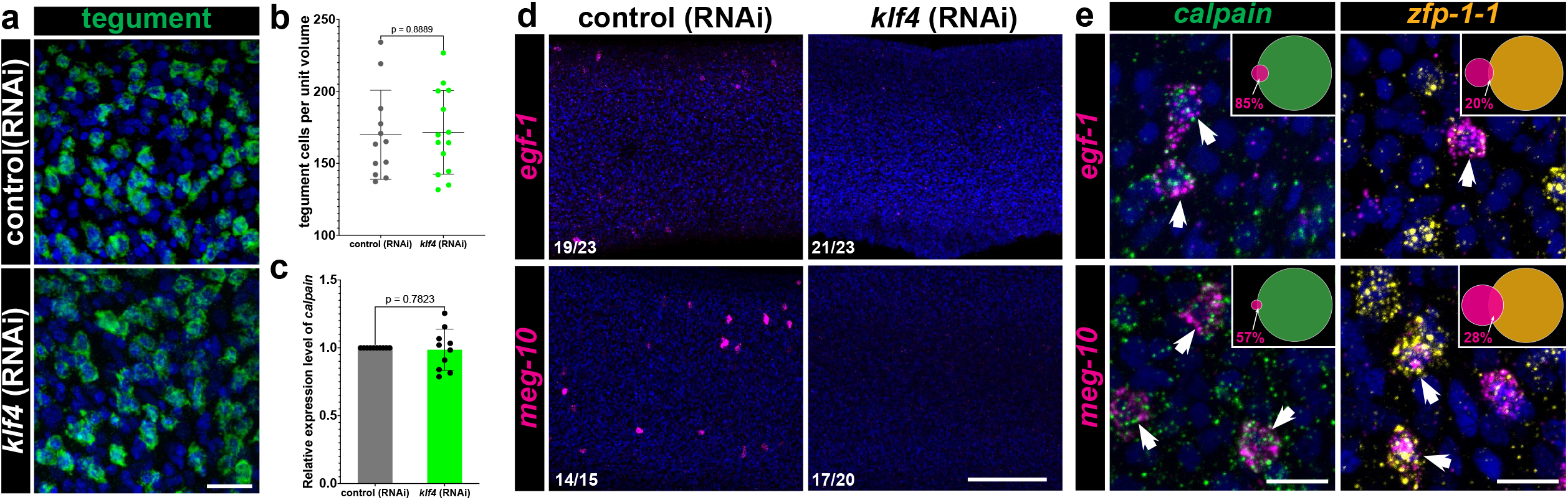
*Klf4*^+^ TAL is required for producing a specific tegumental subpopulation. **a** FISH monitoring the expression for a cocktail of tegumental markers (*calpain, npp-5, annexin* and *gtp-4*) in control (RNAi) and *klf4* (RNAi) worms. Scale bar: 50 µm. **b** Quantification of tegumental cell density. Control (RNAi) n= 12, *klf4* (RNAi) n=14. **c** qPCR detection of expression of the tegumental marker *calpain*. n=10 experiments. **d** FISH results confirm loss of *egf-1*^+^ and *meg-10*^+^ (highlighted with magenta dots in (Fig. 3c)) following *klf4* RNAi. Numbers at bottom left represent the fraction of parasites displaying the observed phenotype. **e** Double FISH showing expression of *egf-1* and *meg-10* relative to the *calpain*^+^ and *zfp-1-1*^+^ cells, respectively (indicated in arrows). The Venn diagram in the upper-right shows the percentage of *egf-1*^*+*^ cells that are *calpain*^+^ cells (201/235 cells, n=17 parasites) or *zfp-1-1*^+^ (12/60 cells n=10 parasites) and the percentage of *meg-10*^*+*^ cells that are *calpain*^+^ or *zfp-1-1*^+^ (127/221 cells are *calpain*^+^, n=13 parasites; 66/233 cells are *zfp-1-1*^+^, n=10 parasites). Scale bars, 100 µm.

### Knockdown of *klf4* alters the heterogeneity of *sm13*^+^ TAL

Since loss of *klf4* resulted in loss of *egf-1*^*+*^ and *meg-10*^*+*^ tegumental cells without compromising total tegumental cell number, we reasoned that there might be a compensatory increase in the rate at which cells in the *sm13*^*+*^ TAL contribute to the tegument. If this were the case, we would anticipate that we would observe an increase in the expression of markers associated with the opposing *sm13*^+^ TAL. Consistent with this prediction, our bulk RNAseq analysis found that of the 12 up-regulated DEGs following *klf4* RNAi (**Fig. 3c, Supplementary Data 2**), 8 were expressed in the *sm13*^+^ TAL (**Supplementary Fig. 4a**). Though *klf4* RNAi did not significantly change the number of *sm13*^*+*^ cells or mRNA levels (**Fig. 5a, 5b**), it led to a 3-fold increase in the expression of *ludp* (*Smp_319910*) (**Fig. 5b**), which encodes a LY6/uPAR domain containing protein, that is predicted to be expressed in *sm13*^+^ cells by scRNAseq (**Supplementary Fig. 4a**). This increase in *ludp* mRNA levels aligns with an observed 2.6-fold increase in the number of *ludp*^*+*^ cells in *klf4* RNAi worms relative to controls (**Fig. 5c left, 5d**). One potential model to explain this increase in *ludp*^+^ cells is that these cells are in a transition state between *sm13*^*+*^ progenitors and *calpain*^*+*^ definitive tegumental cells and loss of *klf4* results in an increase in the flux of cells via this lineage. To test this model, we examined the expression of *ludp* relative to *sm13* and *calpain* in *klf4* (RNAi) parasites. We observed that 14% of *sm13*^+^ cells are *ludp*^+^ in control (RNAi) parasites, and that the proportion of *sm13*^+^ *ludp*^+^ double positive cells jumps to 40% cells in *klf4* (RNAi) worms (**Fig. 5c right, 5e**). We also found that *ludp* was indeed expressed in a small fraction *calpain*^+^ cells (1.6%) and *klf4* RNAi resulted in a significant increase in the number of *ludp*^+^*calpain*^+^ double positive cells (1.6% to 2.4%) (**Fig. 5f, 5g**). These results were mirrored with two other genes (*Lymphocyte antigen 6E-like protein* (*ly6e, Smp_334920*) and *litaf domain containing protein* (*litaf, Smp_333330*) in terms of increases in the number of double positive cells in the *sm13*^+^ TAL **(Supplementary Fig. 4b-d, 4g-I**); however, no increases in the number of double positive cells were noted in the definitive tegumental cell compartment **(Supplementary Fig. 4e-f, j-k)**. These data suggest the possibility that loss of *klf4* prohibits neoblasts from committing to the *egc*/*meg-1*/*zfp-1-1*^+^ TAL, in turn resulting in more *ludp*^+^*sm13*^*+*^ transition state progenitor cells that commit to a *ludp*^+^ tegumental fate. If this were the case, we would anticipate that loss of *klf4*^+^ would result in increased birth of *ludp*^+^*sm13*^+^ transition state cells, yet not comprise the total number of new cells added to the tegument. To test this hypothesis, we knocked down *klf4* and performed an EdU pulse-chase experiment^11^. Specifically, we pulse-labeled neoblasts in control and *klf4* (RNAi) parasites with EdU and monitored the production of EdU^+^ tegumental cells and *ludp*^+^*sm13*^+^ tegument progenitors following a 7D chase. As anticipated, we found *klf4* RNAi led to no change in the percentage of newly produced tegument cells (**Fig. 6a, 6b**). However, we found 52% of *sm13*^+^EdU^+^ cells expressed *ludp*^+^ in *klf4* (RNAi) parasites, which is significantly higher than in control (RNAi) worms (**Fig. 6c, 6d**). Taken together these data suggest that loss of the one TAL leads to an increase in flux through the opposing TAL, in turn altering the molecular make-up of the tegument.

**Fig. 5:**
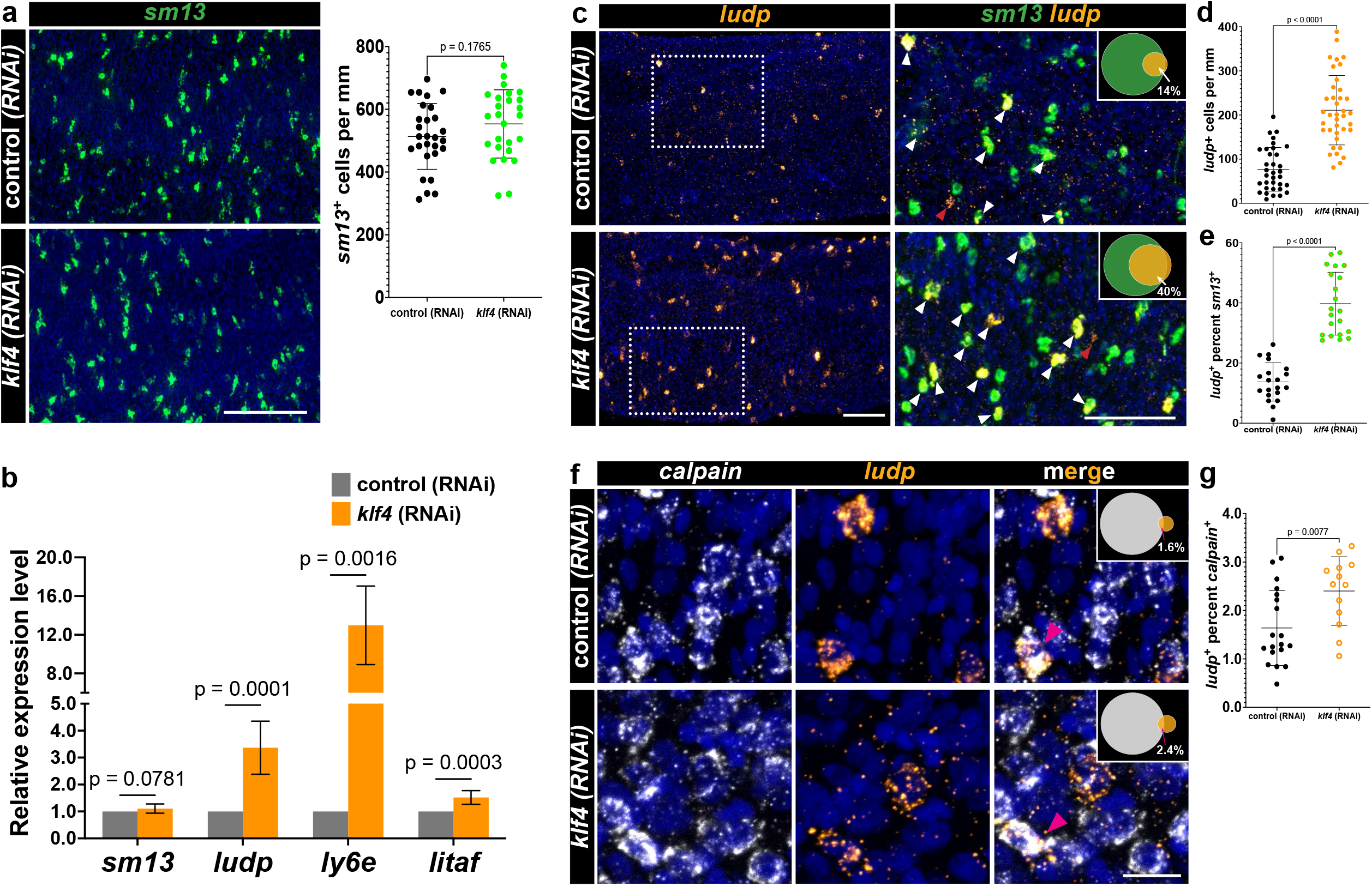
Knockdown of *klf4* alters the heterogeneity of *sm13*^+^ TAL. **a** FISH result showing *klf4* RNAi causes no significant effect on the number of cells expressing *sm13* (left), quantification of the number of *sm13*^+^ cells per mm of worm (right). Control (RNAi) n= 28, *klf4* (RNAi) n=26. **b** qPCR detection of the expression of *sm13* (n=10 experiments) and up-regulated DEGs including *ludp* (*Smp_319910*) (n=10 experiments), *ly6e* (*Smp_334920*) (n=6 experiments) *and litaf* (*Smp_333330*) (n=10 experiments) following *klf4* RNAi. **c** FISH results showing a significant increase in the number of *ludp*^+^ cells following *klf4* RNAi (left); Double FISH showing expression of *sm13* relative to the *ludp*^+^ cells (right); the Venn diagram in upper-right shows the percentage of *sm13*^*+*^ cells expressing *ludp*. White arrows indicate the *sm13*^+^*ludp*^+^ cells, red arrows indicate the *sm13*^-^*ludp*^+^ cells. Scale bar, 100 µm. **d** Quantification of the number of *ludp*^+^ cells per mm of worm. Control (RNAi) n=34, *klf4* (RNAi) n=38. **e** Quantification of percentage of *sm13*^*+*^ cells expressing *ludp*. Control (RNAi) n=20, *klf4* (RNAi) n=21. **f** Double FISH showing expression of *calpain* relative to the *ludp*^+^ cells, and the Venn diagram at up-right shows the percentage of *calpain*^*+*^ cells expressing *ludp*. Control (RNAi) n=19, *klf4* (RNAi) n=13. Scale bar, 10 µm. **g** Quantification of percentage of *calpain*^*+*^ cells expressing *ludp*. Scale bar, 10 µm.

**Fig. 6:**
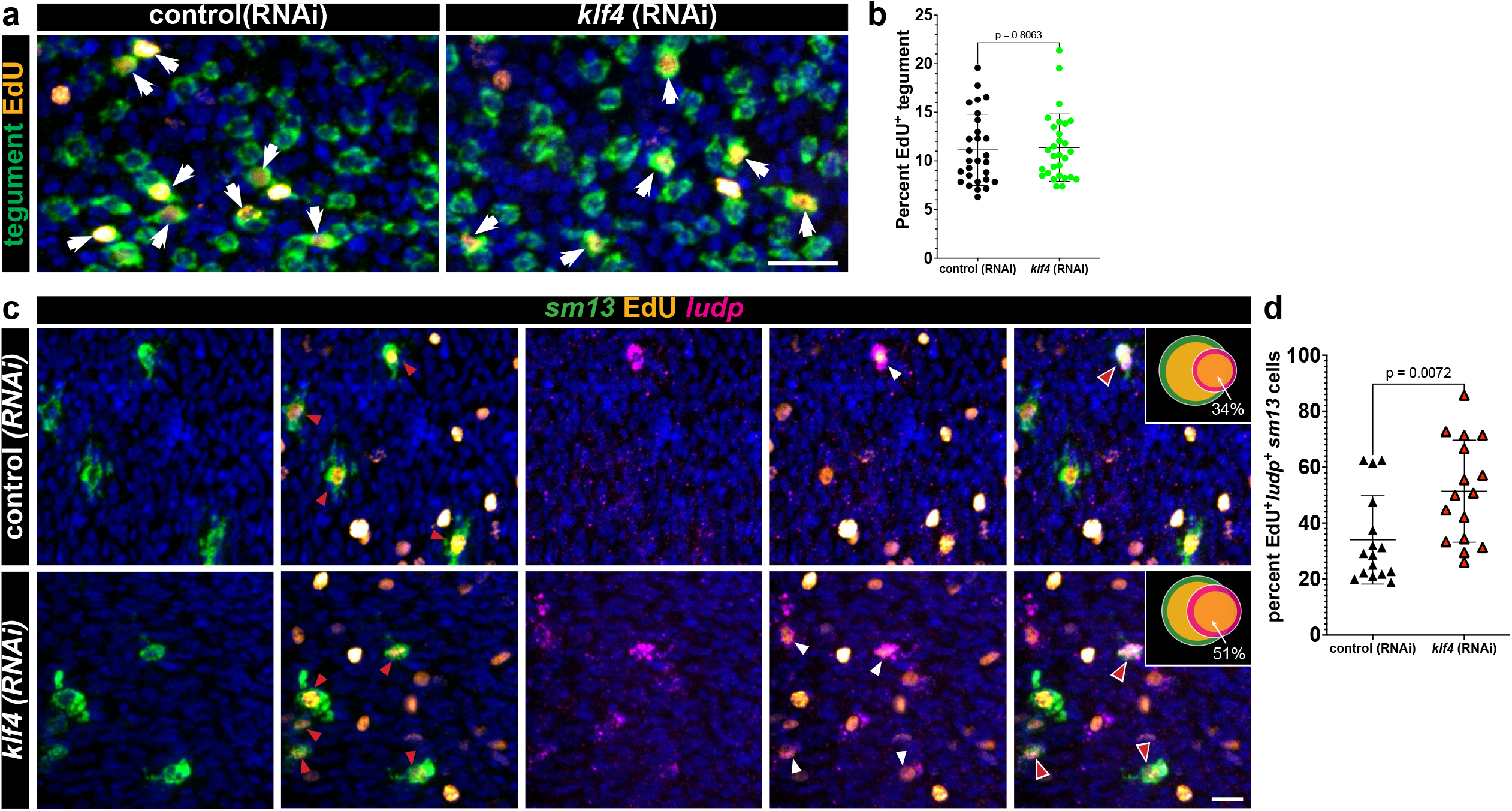
knockdown of *klf4* increases birth of progenitor cells in transition state. **a** FISH for tegumental markers with EdU detection in *klf4* (RNAi) worms at day seven following an EdU pulse. Arrows represent EdU^+^ tegumental cells. **b** Quantification of the percentage of tegumental cells that are EdU^+^ following a 7-day chase period. Control (RNAi) n=28, *klf4* (RNAi) n=29. **c** Double FISH for *sm13* and *ludp* with EdU detection in control (RNAi) and *klf4* (RNAi) worms at day seven following an EdU pulse. Red arrows represent EdU^+^*sm13*^+^ cells, white arrows represent EdU^+^*ludp*^+^ cells and red arrows with white outline represent EdU^+^*ludp*^*+*^*sm13* cells. **d** Quantification of percentage of *ludp*^*+*^*sm13*^*+*^ cells that are EdU^+^ following a 7-day chase period following *klf4* RNAi. Control (RNAi) n=16, *klf4* (RNAi) n=16. Scale bar, 10 µm.

## Discussion

Tegumental maintenance is crucial for schistosome longevity within their human host and depends on sustained turnover from a pool of *tsp-2*^+^ tegument progenitor cells. Here, we reveal that tegumental renewal depends on two opposing *tsp-2*^*+*^ TALs: one that expresses *sm13*^*+*^ and another that expresses *egc*/*meg-1*/*zfp-1-1*^+^. We show that ablation of a Krüppel-like Factor 4 homolog results in loss of the entire *egc*/*meg-1*/*zfp-1-1*^+^ **(Fig. 3b, 3d, 3e)** and loss of *egf-1*^*+*^ tegumental cells **(Fig. 4d)**. In parallel, loss of the *egc*/*meg-1*/*zfp-1-1*^+^ TAL results in an increase in the number of *sm13*^*+*^ and tegumental cells that express *ludp*^+^ **(Fig. 5c-g)**. Surprisingly, while we observe loss of the entire *egc*/*meg-1*/*zfp-1-1*^+^ TAL following *klf4* RNAi treatment we observe no defects in the number of new tegumental cell being born (**Fig. 6a, 6b**). Based on these data we propose a model whereby cells of the *egc*/*meg-1*/*zfp-1-1*^+^ TAL commit to *egf-1*^+^ cells that fuse with the tegument while *sm13*^*+*^ cells commit to *ludp*^+^ cells that likewise fuse with the tegument (**Fig. 7**). Loss of *klf4* blunts commitment of neoblasts to the *egc*/*meg-1*/*zfp-1-1*^+^ TAL resulting in an increase in the rate at which *sm13*^*+*^ lineage seeds new tegumental cell birth. The cumulative effect of *klf4* depletion is an alternation in the molecular composition of the tegumental syncytium.

**Fig. 7:**
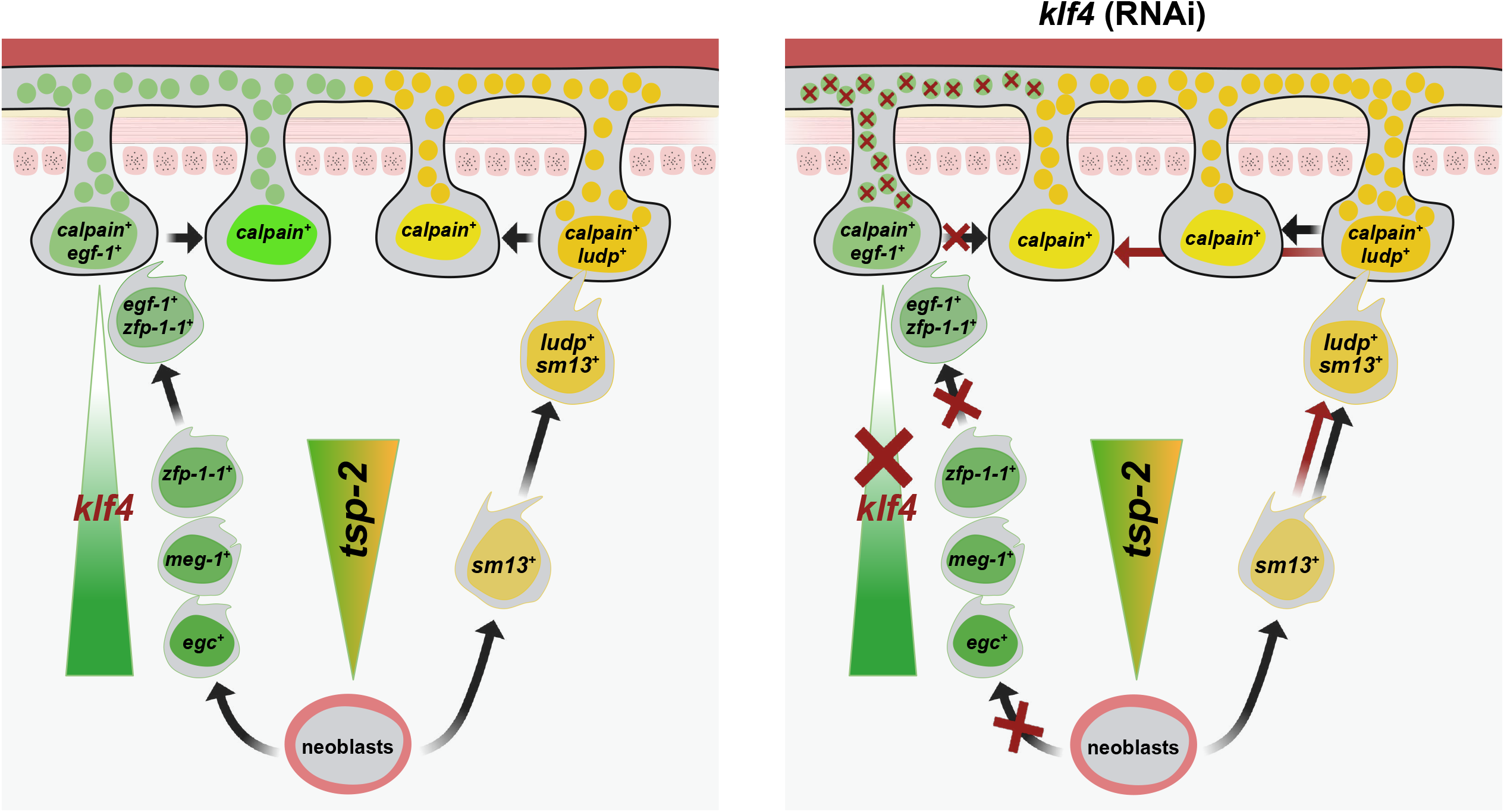
Model of *klf4* involved in tegumental maintenance. Left: The tegument depends on pool of *tsp-2*^+^ tegument progenitor cells. Cells of the *egc*/*meg-1*/*zfp-1-1*^+^ TAL commit to transition state *egf-1*^+^ cells that fuse with the tegument, while *sm13*^*+*^ TAL cells commit to transition state *ludp*^+^ cells that likewise fuse with the tegument. Each TAL delivers a unique set of protein cargos to the tegumental syncytium (green or yellow circles). Right: In *klf4* (RNAi) worms, loss of *klf4* blunts commitment of neoblasts to the *egc*/*meg-1*/*zfp-1-1*^+^ TAL resulting in an increase in the rate at which *sm13*^*+*^ lineage seeds new tegumental cell birth, thereby leading to alternation in the molecular composition of the tegumental syncytium.

Why do the schistosomes appear to specify two molecularly distinct tegumental lineages? Despite being a syncytial tissue with continuous cytoplasm, classic ultrastructural studies of the tegument found various types of tegument-specific cytoplasmic inclusions which appeared to some extent be present in tegumental cells in a mutually-exclusive manner^10,15^. Thus, it is possible that these two lineages are destined to become two specific subtypes of tegumental cells possessing distinct molecular fingerprints and cellular compositions. If this is the case, then why do we only find that only a very small number of *calpain*^*+*^ tegumental cells that express markers indicating the presence of tegumental cell heterogeneity (i.e., *egf-1* and *ludp*) (**Fig. 4d, 5f**)? Given that both *egf-1* and *ludp* are simultaneously expressed in both a subset of *calpain*^*+*^ tegmental cells and subsets of TAL cells, we argue that these are not stable, mature tegumental cells but rather cells that are in a transition state that have recently fused with the tegumental syncytium (**Fig. 7**). Rather than tegumental cells actively expressing cell-type specific mRNAs to establish cellular heterogeneity, we favor a model in which molecularly-distinct tegumental progenitors (i.e., *egc*/*meg-1*/*zfp-1-1*^+^ or *sm13*^*+*^ lineage cells) fuse with the tegument and bring with them a collection of specific proteins to execute distinct molecular functions. Indeed, this model is well-supported as proteins such as TSP-2 and SM13 are known to be present in the tegument^14,16^, yet their mRNA are not observed in mature tegumental cell bodies^11^. Thus, it is likely that many proteins encoded by genes expressed in the respective TALs make their way into the tegument. Understanding specific functions of these TAL-specific genes is expected to illuminate the roles for tegumental subpopulations in parasite biology.

Tegument production and maintenance is a complex process^5^ but pivotal for the success of the parasite in the blood. Here, we uncover *klf4* as a critical transcriptional regulator for maintaining heterogeneity within tegumental cell pool. Although the purpose of this tegumental heterogeneity is unclear, the development of robust experimental models that allow for the ablation of specific TALs in bloodstream schistosomes is anticipated to unravel the functions of the various tegumental cell types. Such studies could reveal the molecular programs that mediate schistosome long-term survival and immune evasion and suggest new therapeutic interventions.

## Materials and methods

### Parasite acquisition and culture

Adult *S. mansoni* (6–7 weeks post-infection) were obtained from infected female mice by hepatic portal vein perfusion with 37°C DMEM (Sigma-Aldrich, St. Louis, MO) plus 10% Serum (either Fetal Calf Serum or Horse Serum) and heparin. Parasites were cultured as previously described (Collins et al., 2016). All experiments were performed with male parasites to maximize the amount of somatic tissue present. Experiments with and care of vertebrate animals were performed in accordance with protocols approved by the Institutional Animal Care and Use Committee (IACUC) of UT Southwestern Medical Center (approval APN: 2017-102092).

### RNA interference

20 freshly perfused male parasites were placed into 6-well plates and cultured in 10mL in Basch Media 169 supplemented with 30 μg/ml dsRNA for 17 days. dsRNA was generated by *in vitro* transcription and replaced with fresh media on Day 0, 1, 2, 5, 8, 11. On day 14, the worms were given fresh media. On day 17, the worms were pulsed with 10 µM EdU for 4 hours before being fixed as previously described^17^; for EdU pulse-chase experiments, the worms were pulsed with 10 µM EdU for 4 hr after which the media was changed, the worms were fixed on day 24 following 7 day chasing period. As a negative control for RNAi experiments, we used a non-specific dsRNA containing two bacterial genes^18^. cDNAs used for RNAi and *in situ* hybridization analyses were cloned as previously described^18^; oligonucleotide primer sequences are listed in **Supplementary Data 5**.

### qPCR and RNAseq analysis

The whole parasites were collected and homogenized in Trizol, RNA extraction, cDNA preparation and qPCR were performed as previously described^19^. Oligonucleotide primer sequences used for qPCR are listed in **Supplementary Data 5**. For RNAseq analysis, three biological replicates were performed for control treatment and knockdown condition. The samples were prepared by Illumina TruSeq stranded mRNA library kit. All 6 samples were sequenced with one flow cell on Illumina NextSeq 550 sequencer with 75bp read lengths. Reads were mapped with STAR (v2.7.10a)^20^ and *S. mansoni* genome sequence (v7) and GTF files used for mapping were acquired from Wormbase Parasite^21^. Differential gene expression were performed with DESeq2 (version 1.30.1)^22^ R(4.0.3). Raw and processed data have been deposited in NCBI (GSE268037). Volcano plots were made with plotting Log2 fold change expression and -Log10 (*P*adj) of differential expressed genes (*P*adj < 0.05) in GraphPad Prism.

### Fluorescence activated cell sorting

FACS sorting was performed as previously described^13^ with minor modifications. After RNAi experiments, 100 male worms from each group were suspended in a 0.5% solution of Trypsin/EDTA (Sigma T4174) in PBS before rinsing in PBS for triple times. The worms were then triturated for approximately 10 minutes at room temperature until the solution became turbid and no large pieces of worms were left. The trypsin was inactivated by adding an equal volume of Basch media. The dissociated worms were then centrifuged at 500 g for 10 min at 4°C. Next the worms were resuspended in 1 ml of Basch medium followed by filtering through 100uM cell strainer to remove big chunks. The filtered collections were treated with 10 μL of RQ1 DNAse (Promega M6101) and incubated for 10 minutes at RT. After adding 3mL PBS, the dissociated worms were centrifuged again at 500 g for 10 minutes at 4°C. The cells were then resuspended in 1mL of staining media (0.2% BSA, 2mM EDTA in PBS, pH 7.40) and incubated in Hoechst 33342 (18 μg/ml) (Sigma B2261) for 30min at RT in the dark. The worms were centrifuged once again at 500 g for 10 minutes at 4°C. Worms were then resuspended in 1 mL of staining media containing Hoechst 33342 (18 μg/ml) and propidium iodide (1 μg/ml) (Sigma-Aldrich P4170) and then filtered through a 100 μm cell strainer into a 12×75mm FACS tube prior to sorting. Filtered cells were then sorted on a FACSAria II custom (BD Biosystems) with 305/405/488/561/633nm lasers. Live single cells (PI negative, singlet by comparing forward scatter height to forward scatter width) were sorted using a 100 μm nozzle and cells were sorted into sorting media (0.2% BSA in PBS, pH 7.4). A Hoechst threshold was applied to exclude debris and improve the efficiency of sorting.

### Single-cell RNA sequencing

FACS-sorted cells were centrifuged again at 500 g for 10 minutes at 4°C then resuspended in 0.2% BSA in PBS. Libraries were created using a Chromium Controller (10x Genomics) according to manufacturer guidelines and sequenced in using a NextSeq 500 (illumina). Sequencing data was processed and mapped to the *Schistosoma mansoni* genome (v7) using Cell Ranger (10x Genomics). Unfiltered data from Cell Ranger was imported into Seurat (v4.3)^23,24^ and cells were filtered as follows: nFeature_RNA > 200 & nFeature_RNA<4000 & nCount_RNA > 1000 & nCount_RNA < 20000 & percent.mt < 5 (Mitochondrial genes were identified as those with the prefix “Smp_9”). Control (RNAi) and *klf4* (RNAi) datasets was normalized (NormalizeData) and variable features were identified (FindVariableFeatures, selection.method = “vst”, nfeatures =2000). From here, integration anchors were identified (FindIntegrationAnchors, dims 1:60), the data was integrated (IntegrateData, dims = 1:60, features.to.integrate = all.genes), and scaled (ScaleData). We then ran RunPCA, RunUMAP (reduction = “pca”, dims = 1:60, n.neighbors =30), FindNeighbors (reduction = “pca”, dims = 1:60), FindClusters (resolution = 4). After merging we were left with a final map of 65 clusters of 12910 cells. To identify the clusters of tissue distribution on the UMAP, we used FindAllMarkers (avg_log2FC>1) command to identify marker genes of each cluster (**Supplementary Data 3**), and validated by manually inspecting tissue markers, including *egc*^*+*^(*Smp_314170*), *meg-1*^*+*^(*Smp_122630*), *zfp-1-1*^*+*^ (*Smp_049580*), early *tsp-2*^+^ (*Smp_335630*), *sm13*^+^ (*Smp_195190*), tegument (*calpain, Smp_214190*), tegument2 (*Smp_056460*), neoblast progeny (*Smp_171720; hes2 Smp_132810*), neoblasts (*nanos2 Smp_051920; eled Smp_041540*), neurons (*7b2, Smp_073270*), flame cells (*sialidase Smp_335600*), muscle (*tpm2, Smp_031770*), parenchyma (*tgfbi, Smp_212710*) and gut (*hnf4 Smp_174700*; *ctsb, Smp_103610*)^13^. Then we collapsed all tissue-specific clusters into a single cluster and assigned them same color and name, generated the UMAP of merged datasets (**Supplementary Figure 2**) and the marker list of each labeled cluster (**Supplementary Data 4**). The dot plots for Fig.3d, Supplementary Figs.2f and 4a were generated using the DotPlot() function in Seurat v4.3 with the all down-regulated genes and up-regulated genes following *klf4* RNAi (**Supplementary Data 2**). The size of the dot corresponds to the percentage of the cells in the cluster (indicated on the vertical axis) that express the given gene (indicated on the horizontal axis), whereas the color of the dot indicates the average expression level of the gene in the cluster. To identify differential expression genes in same cell types between control (RNAi) and *klf4* (RNAi) worms, since *egc*^*+*^ and *meg-1*^*+*^& *zfp-1-1*^*+*^ populations were too small to perform differential analysis, we combined the *egc*^*+*^, *meg-1*^*+*^& *zfp-1-1*^*+*^, tegument and tegument2 clusters into a single “*klf4*^+^ TAL” cluster and combined the early *tsp-2*^+^ and *sm13*^+^ clusters into a single “*sm13*^+^ TAL” cluster, then using the FindMarkers() command to identify differential expressed genes of these clusters (**Supplementary Data 1**). Raw and processed data have been deposited in NCBI (GSE268036).

### Parasite labeling and imaging

Colorimetric and fluorescence *in situ* hybridization detections were performed as previously described^12,17^ with the following modification. To improve signal-to-noise for colorimetric *in situ* hybridization, all probes were used at 10 ng/mL in hybridization buffer. For FISH, we prolonged the TSA reaction time to 20 or 40min with 50 ng/mL probe or 10ng/mL probe, respectively. EdU detection was performed as previously described^17^. All fluorescently labeled parasites were counterstained with DAPI (1 μg/mL), cleared in 80% glycerol, and mounted on slides with Vectashield (Vector Laboratories).

Confocal imaging of fluorescently labeled samples was performed on either a Zeiss LSM900 or a Nikon A1 Laser Scanning Confocal Microscope. To perform cell counts, cells were manually counted in maximum intensity projections derived from confocal stacks. In cases where we determined the number of cells in a particular region of the parasite (e.g., tegument) we collected confocal stacks and normalized the number of cells by total volume of the stack in μm^3^. In cases where we determined the total number of labeled foci throughout the entire depth of the parasite (e.g. *tsp-2*^*+*^ counts), we collected confocal stacks and normalized the number of cells to the length of the parasite in the imaged region in mm. The Venn diagrams were generated by https://academo.org/demos/venn-diagram-generator/ using the percentage of each interactive cell populations, calculated from cell counts. Brightfield images were acquired on a Zeiss AxioZoom V16 equipped with a transmitted light base and a Zeiss AxioCam 105 Color camera.

### Statistical analysis

GraphPad Prism software processed and presented the data as the mean with SD. All two-way comparisons were analyzed using Welch’s t-test, and multiple paired t-tests were performed in Fig. 5b and Supplementary Figs. 2g and 3a.

## Acknowledgements

The Schistosome Infected mice and B. glabrata snails were provided by the National Institute of Allergy and Infectious Diseases (NIAID) Schistosomiasis Resource Center of the Biomedical Research Institute (Rockville, MD, USA) through National Institutes of Health (NIH)-NIAID Contract HHSN272201700014I for distribution through BEI Resources. FACS was performed with the aid of the Moody Foundation Flow Cytometry Facility at the University of Texas Southwestern Medical Center (UTSW). RNAseq was performed with the aid of the Genomics Sequencing Core at UTSW. Biorender was used for some schematic material. This work was supported by the National Institutes of Health R01AI121037 (J.J.C.) and R01AI150776(J.J.C.), as well as the Welch Foundation I-1948-20240404 (J.J.C.).

## Author contributions

L.Z. performed all the experiments, collected and analyzed data, G.R.W contributed to the conception and design of the research and cell sorting experiment, J.J.C supervised the project and provided conceptual advices, L.Z. drafted the manuscript with revisions from G.R.W and J.J.C.

