## Supplementary material for "A Krüppel-like factor establishes cellular heterogeneity during schistosome tegumental maintenance": Sup. Figures

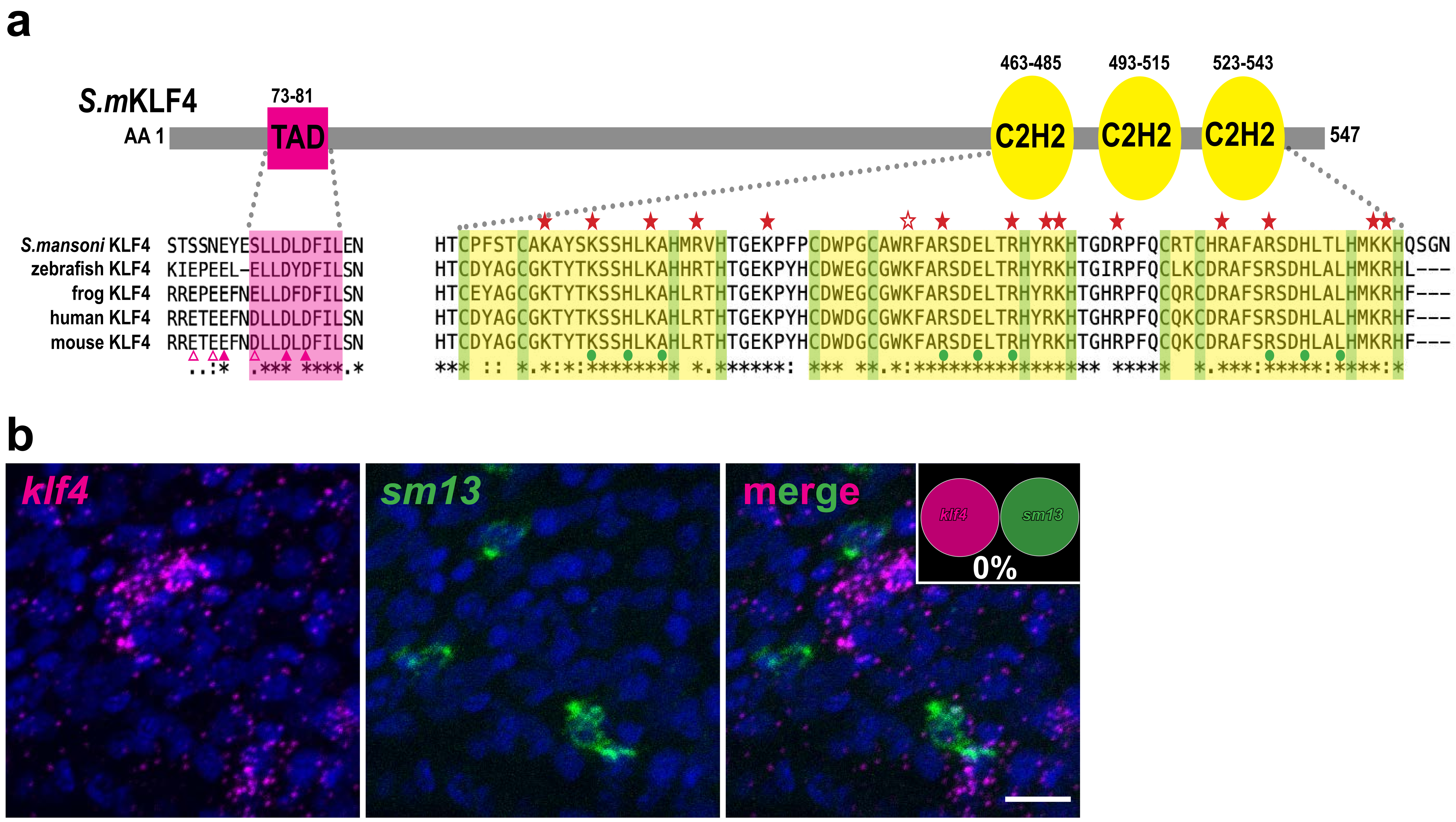


**Supplementary Figure 1:**

**a** Protein sequence alignment of *Sm*KLF4 with KLF4 from *Danio rerio* (NP_001106955.1), *Xenopus tropicalis* (NP_001017280.1), *Mus musculus* (NP_034767.2) and *Homo sapiens* (NP_001300981.1). *Sm*KLF4 is 547 amino acids in length, and contains a 9AA transactivation domain (TAD) located at position AA73-81 (highlighted in pink rectangle). The acidic amino acids glutamine (E) and aspartate (D) in humans are indicated with pink triangles, filled triangle represents identical residue in the *S. mansoni* protein and empty triangles indicates non-identical residues. At the C-terminus, *Sm*KLF4 features an 81AA highly conserved region containing three C2H2 zinc finger domains. The amino acid sequence below is highlighted in filled yellow rectangles, with cystine and histidine residues highlighted in green. Fifteen highly conserved basic residues (KKKRKr/kRRRK RRRKK) are marked with red stars, with only one non-identical residues labelled with an empty star, nine conserved residues crucial for DNA binding specificity are highlighted with filled green circles. **b** *klf4* (246 cells) has no expression in tegument progenitor *sm13*$^{+}$ cells (250 cells) (0/250, n=6 parasites). Scale bar, 10 μM.

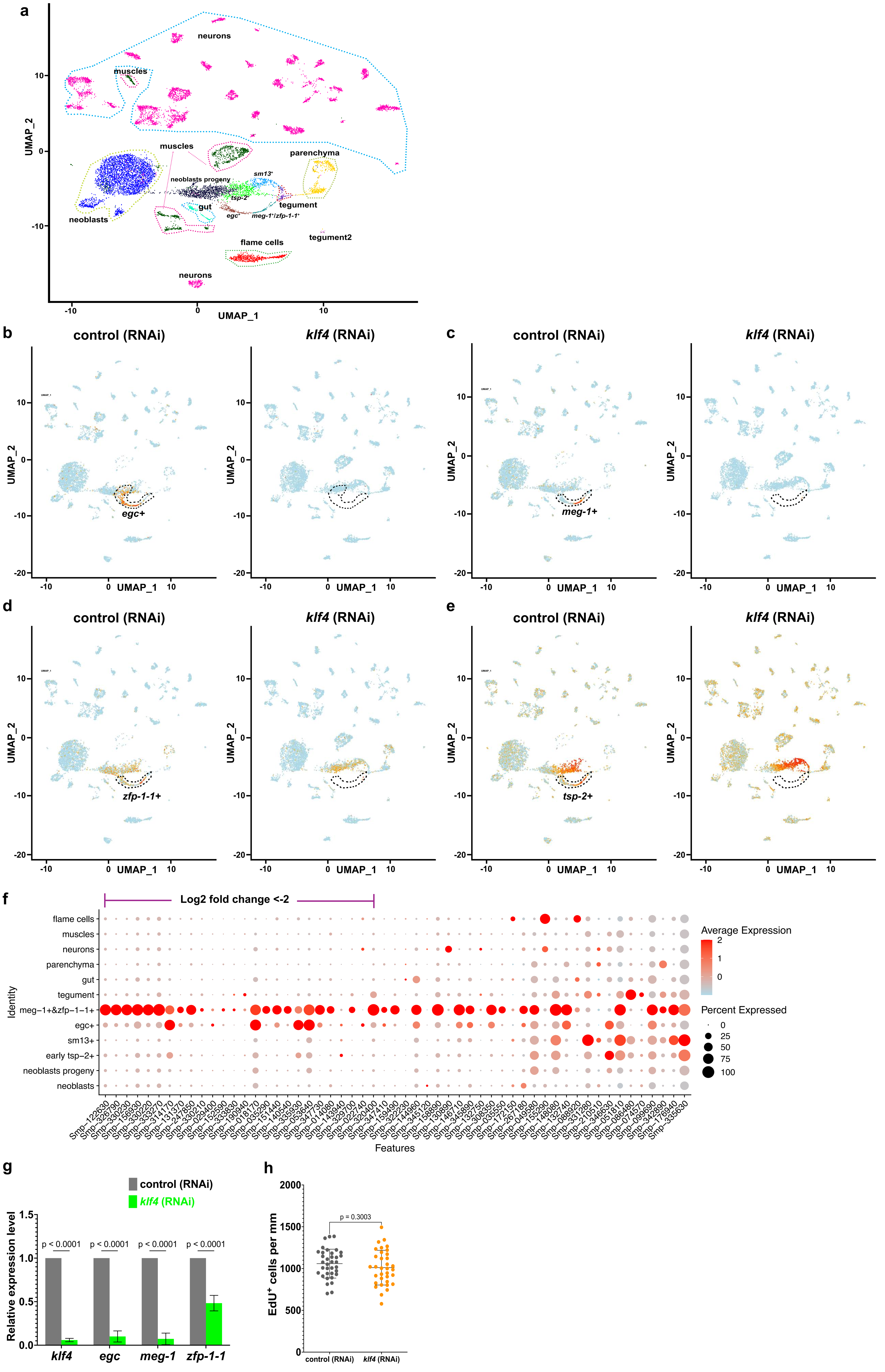

**Supplementary Figure 2:**
**a** UMAP plot of all clusters and their predicted cellular identity from *klf4* RNAi scRNAseq analysis. **b-e** UMAP plot shows expression of (**b**) *egc*, (**c**) *meg-1*, (**d**) *zfp-1-1* and (**e**) *tsp-2* in control (RNAi) and *klf4* (RNAi) worms. **f** A dot-plot summarizing the expression of all bulk RNAseq down-regulated DEGs in the *klf4* RNAi-mediated scRNAseq profile. Cluster populations are on the vertical axis and gene IDs are on the horizontal axis. Expression levels are colored by gene expression (blue = low, red = high). Percentage of cells in the cluster expressing the gene is indicated by the size of the circle. **g** qPCR detection of expression of *klf4* (n=10 experiments), *egc* (n=8 experiments), *meg-1* (n=7 experiments) and *zfp-1-1* (n=9 experiments) following *klf4* RNAi. **h** Quantification of the number of EdU⁺ per mm of worms. Control (RNAi) n= 35, *klf4* (RNAi) n=34.

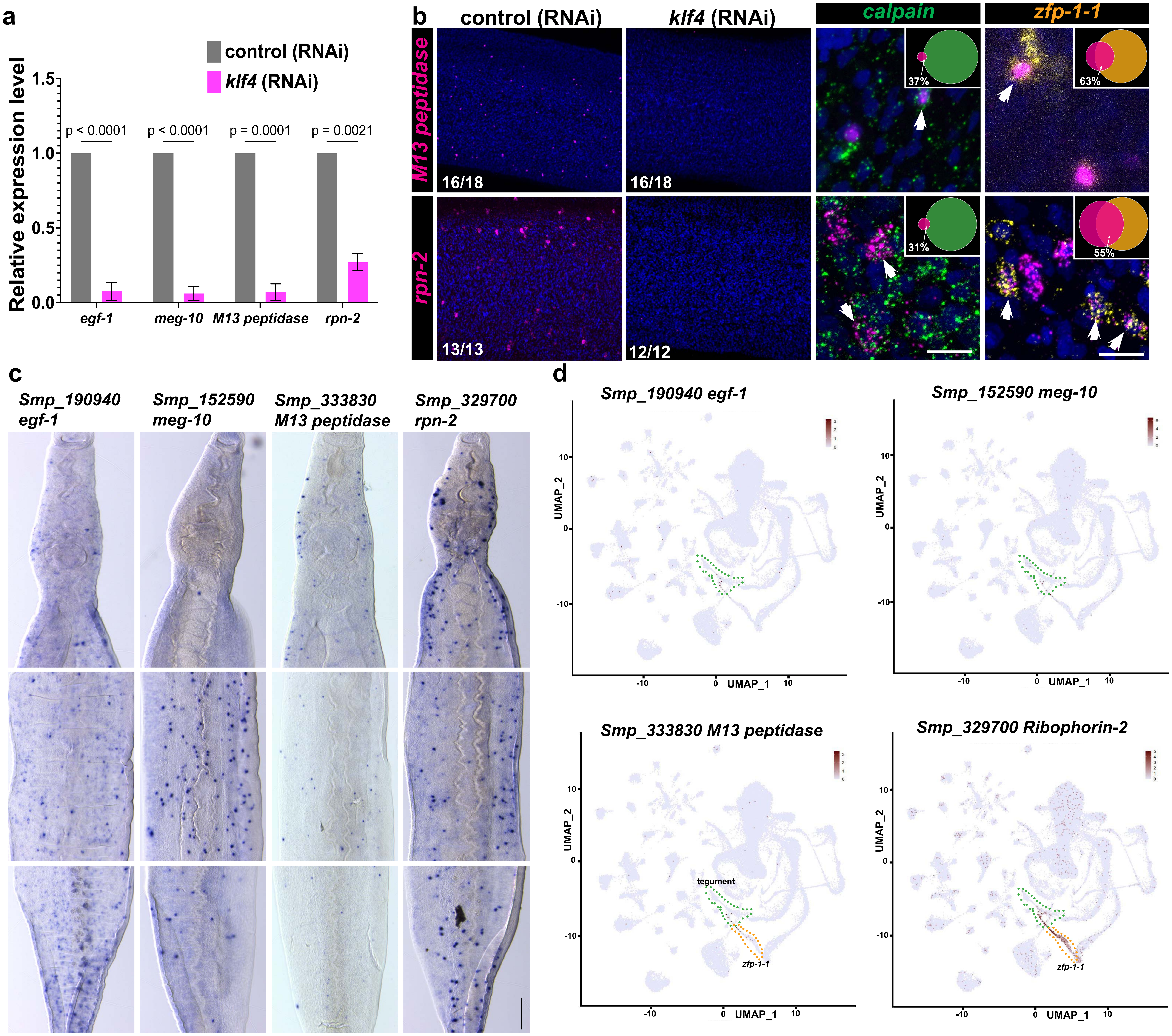


**Supplementary Figure 3:**

**a** qPCR dtection of expression of *egf-1* (n=5 experiments), *meg-10* (n=6 experiments), *M13 peptidase* (n=4 experiments) and *rpn-2* (n=3 experiments) following *klf4* RNAi. **b** Left, FISH results confirm a loss of *M13 peptidase*$^+$ and *rpn-2*$^+$ cells (highlight with magenta dot in (Fig. 3c)) following *klf4* RNAi. Numbers at bottom left represent the fraction of parasites displaying the observed phenotype. Right, Double FISH showing expression of *M13 peptidase*$^+$ and *rpn-2*$^+$ relative to the *calpain*$^+$ and *zfp-1-1*$^+$ cells, respectively (indicated in arrows). The Venn diagram in upper right shows the percentage of *M13 peptidase*$^+$ (114/310 cells are *calpain*$^+$, n=16 parasites; 45/71 cells are *zfp-1-1*$^+$ , n=6 parasites) and *rpn-2*$^+$ (197/637 cells are *calpain*$^+$, n=10 parasites; 212/384 cells are *zfp-1-1*$^+$, n=9 parasites) in *calpain*$^+$ and *zfp-1-1*$^+$ cells. Scale bar, 100 μm. **c** WISH showing expression pattern of *egf-1*, *meg-10*, *M13 peptidase* and *rpn-2*. Scale bar, 100 μM. **d** UMAP plot shows expression pattern of *egf-1*, *meg-10*, *M13 peptidase* and *rpn-2* on the established adult scRNAseq atlas.

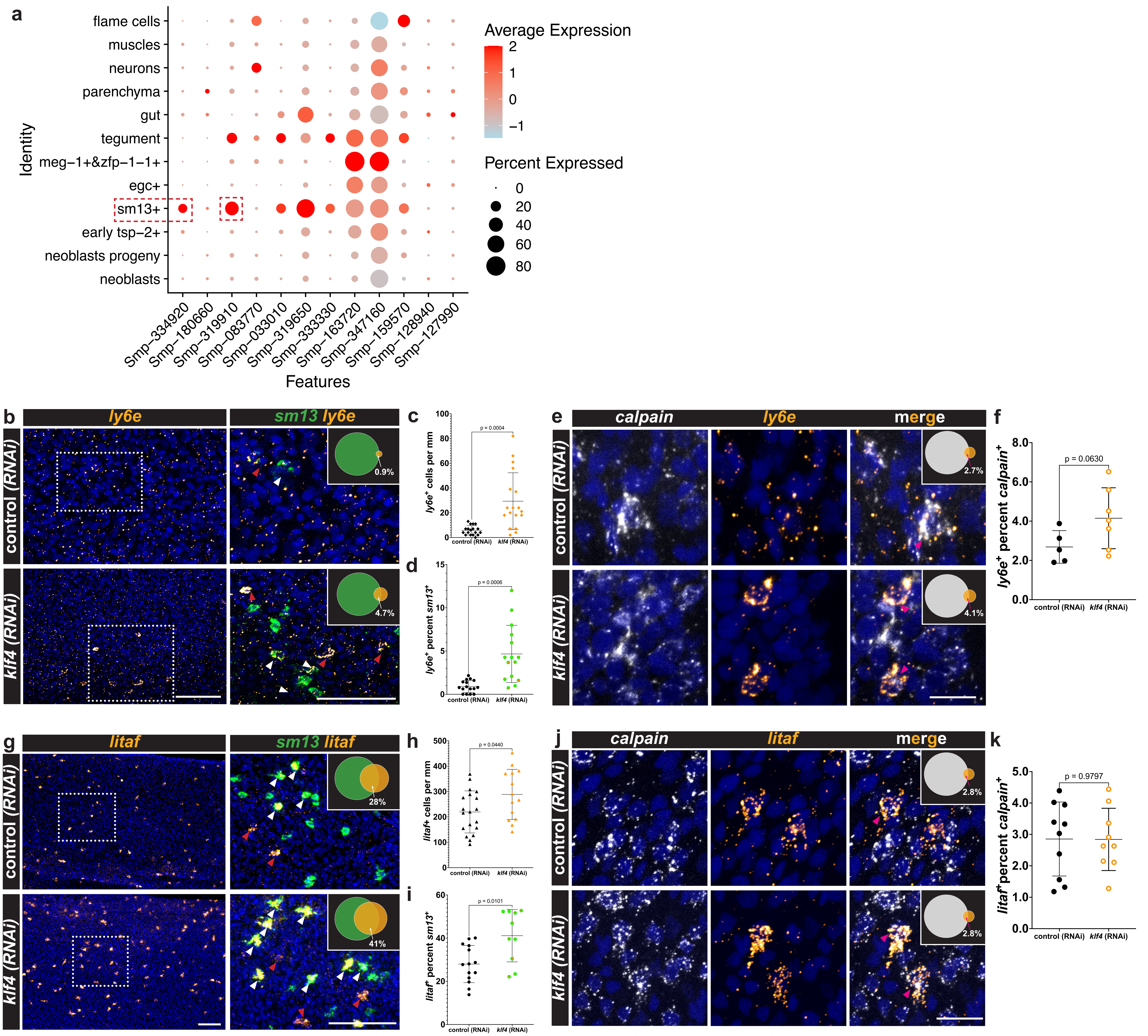


**Supplementary Figure 4:**

**a** Dot-plot summarizing the expression of the up-regulated DEGs from the bulk RNAseq analysis on the *klf4* RNAi scRNAseq profile. The genes enriched in the *sm13*⁺ cell population are indicated in dashed rectangle. **b** FISH results showing an increase in the number of *ly6e*⁺ cells following *klf4* RNAi (left); double FISH showing expression of *sm13* relative to the *ly6e*⁺ cells (right), white arrows indicate the *sm13*⁺*ly6e*⁺ cells and red arrows indicate the *sm13*⁻*ly6e*⁺ cells. Venn diagram in upper right shows the percentage of *sm13*⁺ cells expressing *ly6e*. Scale bar, 100 μm. **c** Quantification of the number of *ly6e*⁺ cells per mm of worms. Control (RNAi) n=18, *klf4* (RNAi) n=18. **d** Quantification of percentage of *sm13*⁺ cells expressing *ly6e*. Control (RNAi) n=16, *klf4* (RNAi) n=15. **e** Double FISH showing expression of *calpain* relative to the *ly6e*⁺ cells, pink arrows indicate the *calpain*⁺*ly6e*⁺ cells. Venn diagram in upper right shows the percentage of *calpain*⁺ cells expressing *ly6e*. **f** Quantification of the percentage of *calpain*⁺ cells expressing *ly6e*. Control (RNAi) n=5, *klf4* (RNAi) n=7. **g** FISH results showing an increase in the number of *litaf*⁺ cells following *klf4* RNAi (left); double FISH showing expression of *sm13* relative to the *litaf*⁺ cells (right), white arrows indicate the *sm13*⁺*litaf*⁺ cells, red arrows indicate the *sm13*⁻*litaf*⁺ cells. Venn diagram in upper right shows the percentage of *sm13*⁺ cells expressing *litaf*. Scale bar, 100 μm. **h** Quantification of the number of *litaf*⁺ cells per mm of worms. Control (RNAi) n=19, *klf4* (RNAi) n=14. **i** Quantification of percentage of *sm13*⁺ cells expressing *litaf*. Control (RNAi) n=14, *klf4* (RNAi) n=10. **j** Double FISH showing expression of *calpain* relative to the *litaf*⁺ cells, and the percentage of *calpain*⁺ cells expressing *litaf* gene was quantified that shown in Venn diagram in upper right. **k** Quantification of percentage of *calpain*⁺ cells expressing *litaf*. Control (RNAi) n=10, *klf4* (RNAi) n=9. Scale bar, 10 μm.

### Description of Supplementary Data

**Supplementary Data 1:** Differentially expressed analysis within the same cell type from single cell-RNAseq (scRNAseq) datasets between control (RNAi) and *klf4* (RNAi) conditions.

**Supplementary Data 2:** Differentially expressed genes (DEGs) from RNAseq analysis following *klf4* RNAi.

**Supplementary Data 3:** List of genes significantly enriched in every single cluster (65 clusters in total) from scRNAseq analysis.

**Supplementary Data 4:** List of genes significantly enriched in the same cell types from scRNAseq analysis.

**Supplementary Data 5:** List of oligos and abbreviations used in this study.
